# Mining Pathway Associations from Networks of Mutual Exclusivity Interactions

**DOI:** 10.1101/2020.02.20.957241

**Authors:** Herty Liany, Yu Lin, Anand Jeyasekharan, Vaibhav Rajan

## Abstract

Study of pairwise genetic interactions such as mutual exclusivity or synthetic lethality has led to the development of targeted anticancer therapies, and mining the network of such interactions is a common approach used to obtain deeper insights into the mechanism of cancer. A number of useful graph clustering-based tools exist to mine interaction networks. These tools find subgraphs or groups of genes wherein each gene belongs to a single subgraph. However, a gene may be present in multiple groups – for instance, a gene can be involved in multiple signalling pathways. We develop a new network mining algorithm, that does not impose this constraint and can provide a novel pathway-centric view. Our approach is based on finding edge-disjoint bipartite subgraphs of highest weights in an input network of genes, where edge weights indicate the significance of the interaction and each set of nodes in every bipartite subgraph is constrained to belong to a single pathway. This problem is NP-hard and we develop an Integer Linear Program to solve this problem. We evaluate our algorithm on breast and stomach cancer data. Our algorithm mines dense between-pathway interactions that are known to play important roles in cancer and are therapeutically actionable. Our algorithm complements existing network mining tools and can be useful to study the mutational landscape of cancer and inform therapy development.

## 1 Introduction

Cancer cells gain selective advantage over healthy cells and proliferate uncontrollably as a result of genetic alterations caused by endogenous and exogenous processes [23]. These alterations are observed in multiple genes that participate in important signalling pathways, and the mechanisms of these alterations differ across cancer types and subtypes [4, 39]. Investigation of combinatorial patterns of alterations, in genes and pathways, have enabled us to understand cancer better and find effective therapeutic targets.

Genetic alterations affecting pairwise interactions, such as Mutually Exclusive (ME) mutations have been studied extensively. ME can indicate functional redundancy or Synthetic Lethality (SL) [17]. SL interactions between two genes is a condition where the loss of either gene is viable but the loss of both is lethal, and has been considered a foundation for development of targeted anti-cancer therapies [7, 40]. Such interactions across pathways can result in cross-talk between pathways [30, 46] and methods have been developed to identify such between-pathway motifs [6, 24]. Many other computational approaches have also been developed to infer ME [12, 3, 32, 27, 13, 8] and SL interactions [28, 35, 45, 43, 31, 33].

A network provides a succinct global representation of all the pairwise genetic interactions observed in a set of genes. Analysis and mining of such networks can not only yield important biological insights but have also aided the development of predictive models for gene essentiality, survival and drug response [28, 31]. A common approach for exploratory mining is to find biologically interesting sub-networks, also called modules. Recent examples of methods to find modules include BeWith [15], CDPath [56] and CD-CAP [25].

BeWith [15] identifies modules with combinations of interactions both within and between modules. Their clustering approach is formulated as an Integer Linear Program (ILP) to optimize between-cluster and within-cluster scores. The scores may be based on functional relatedness, mutual exclusivity or co-occurrence. CDPath [56] uses an ILP similar to that of BeWith to identify modules that maximize coverage and mutual exclusivity within modules, and co-occurrence and functional interactions between modules. Pathways associated with each module are identified using gene-pathway association information. Further, it clusters the pathway interaction network to group pathways with functional relations. Pathways in different modules and same clusters are considered as cooperative driver pathways. Combinatorial Detection of Conserved Alteration Patterns (CD-CAP) [25] detects subnetworks of an interaction network each with an alteration pattern, within a group of genes, conserved across tumor samples. The patterns include copy number alterations, somatic mutations and aberrant gene expression. Although these methods differ in their approaches and objectives, they can all be viewed as variants or extensions of graph clustering methods.

In this paper we develop another approach to mine and analyse a given network of pairwise interactions, that complements existing approaches. While graph clustering is useful to find modules, the standard cut-based approach only finds groups of genes wherein each gene belongs to a single sub-network. They cannot find patterns of interactions across groups where a gene may be present in multiple groups, which is biologically plausible – for instance, a gene can be involved in multiple signalling pathways.

A schematic of our approach is shown in figure 1. Our approach is based on finding *K* edge-disjoint bipartite subgraphs of highest weights in an input network of genes, with edge weight indicating the significance of the interaction. Further, we constrain each edge set of the bipartite subgraphs to belong to a single pathway. Thus, each bipartite subgraph in our solution represents a dense between-pathway interaction. While each interaction can only be part of a single subgraph, there are no restrictions on the genes that may belong to multiple subgraphs. This problem is NP-hard and we develop an ILP-based heuristic to solve this problem.

**Fig. 1.**
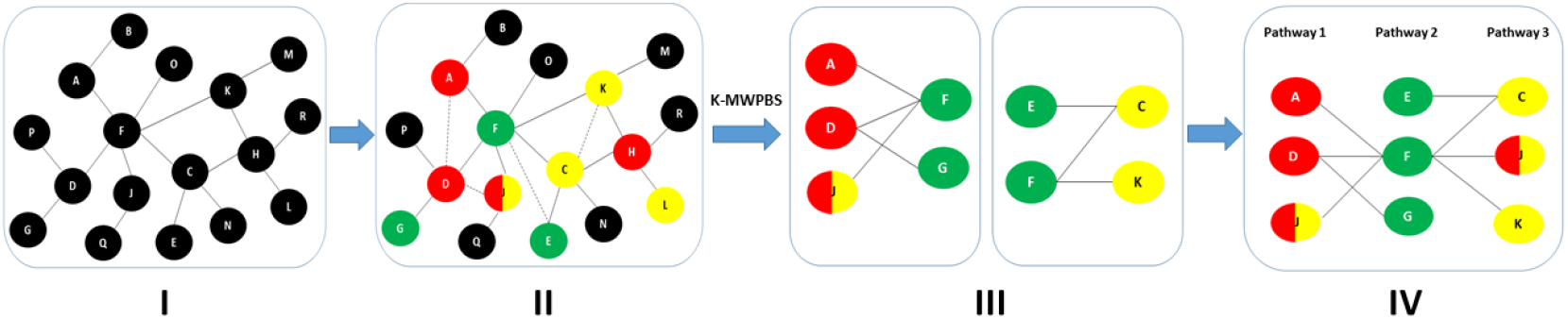
(I) Input network. Nodes are genes, edges are pairwise interactions, e.g. ME, SL. Edge weights not shown. (II) Same network in I with information of input pathways, in which some of the genes participate; each color (green, red, yellow) indicates a pathway. A gene may belong to more than one pathway, e.g. nodes with 2 colors are in 2 pathways. Dashed lines indicate within-pathway edges. Genes that are not involved in any of the input pathways are in black. (III) The bipartite subgraphs obtained by our algorithm using input network and pathway information from II. Each bipartite subgraph shows the interactions between gene sets from two different pathways (left: red-green, right: green-yellow). Within-pathway edges are not considered. (IV) The bipartite subgraphs in III merged to provide a pathway-centric view. Our results in later sections are shown in this format.

We use our approach to explore ME interaction networks in breast and stomach cancers. Our algorithm uncovers relationships between signalling pathways that are known to play important roles in cancer and in targeted drug development. Such between-pathway relationships can potentially help elucidate functional mechanisms of oncogenic alterations [39]. We believe that our tool complements the existing repertoire of network mining tools that aim to gain deeper insights into the mutational landscape of cancer and inform therapy development.

## 2 Method

The inputs to our method are (1) an undirected graph *G* = (*V, E*) with vertex set *V* and weighted edge set *E*, and (2) a list of pathways *P*, where a pathway *P*_*j*_ is represented by a list of participating genes. In our context, a vertex represents a gene and the edges *E* represent ME relations.

A weighted bipartite graph, (*L, R, F*), is a graph whose vertices can be divided into disjoint sets, denoted by left (*L*) and right (*R*) sets, and every weighted edge in the edge set *F* is such that it connects a vertex in *L* to a vertex in *R*. The edges, *F*, in a bipartite graph show us the interactions between (the left and right) sets of genes. We define the weight, *w*(*F*), of a bipartite graph as the sum of the weights of all the edges in the graph. We require every gene in the left set of the mined bipartite graph to belong to one of the input pathways, that we call the *left pathway*. Similarly, every gene in the right set of the mined bipartite graph is required to belong to one of the input pathways, that we call the *right pathway*. To find those pathway pairs with the most ‘dense’ interactions, we mine bipartite subgraphs of *G* with maximum weight. Note that we do not pose any restrictions on the edges between genes within a pathway. Thus, the input network may have within-pathway ME interactions, that are not shown in the subgraphs. These requirements are modeled through the following combinatorial optimization problems.

The *Maximum Weight Pathway Constrained Bipartite Subgraph* problem (MWPBS) is to find a bipartite subgraph (*L, R, F*) of an input graph (*V, E*) that is of maximum weight and such that every gene in the left and right sets of the subgraph belongs to a pathway and the associated left and right pathways are different, i.e.,

1. *L* ⊂ *V*, *R* ⊂ *V*, *L* ∩ *R* = *ϕ*, *F* = {(*u, v*)|*u* ∈ *L*, *v* ∈ *R*} ⊆ *E*
2. *L* ⊆ *P*_*k*_, *R* ⊆ *P*_*l*_, *P*_*k*_ ≠ *P*_*l*_
3. *w*(*F*) is maximum among all such subgraphs (satisfying conditions 1 and 2).

The *Top-K Maximum Weight Pathway Constrained Bipartite Subgraph* Problem (K-MWPBS) is to find K edge-disjoint maximum weight pathway constrained bipartite subgraphs of the input graph, (*L*_1_*, R*_1_*, F*_1_), …, (*L*_*K*_, *R*_*K*_, *F*_*K*_), i.e.,

1. *L*_*i*_ ⊂ *V*, *R*_*i*_ ⊂ *V*, *L*_*i*_ ∩ *R*_*i*_ = *ϕ*, *F*_*i*_ = {(*u, v*)|*u* ∈ *L*_*i*_, *v* ∈ *R*_*i*_} ⊆ *E*, ∀*i* ∈ [*K*]
2. *F*_*i*_ ∩ *F*_*j*_ = *ϕ* ∀*i, j* ∈ [*K*]
3. *L*_*i*_ ⊆ *P*_*k*_, *R*_*j*_ ⊆ *P*_*l*_, *P*_*k*_ ≠ *P*_*l*_
4. *w*(*F*_1_) ≥ *w*(*F*_2_) ≥ … ≥ *w*(*F*_*K*_) such that 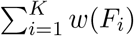 is maximum among all such K subgraphs (satisfying conditions 1–3).

Given a procedure to solve the MWPBS problem, we can iteratively solve the K-MWPBS problem in a greedy manner as shown in algorithm 1. Note that the subgraphs are edge disjoint but may share vertices.

**Algorithm 1:**
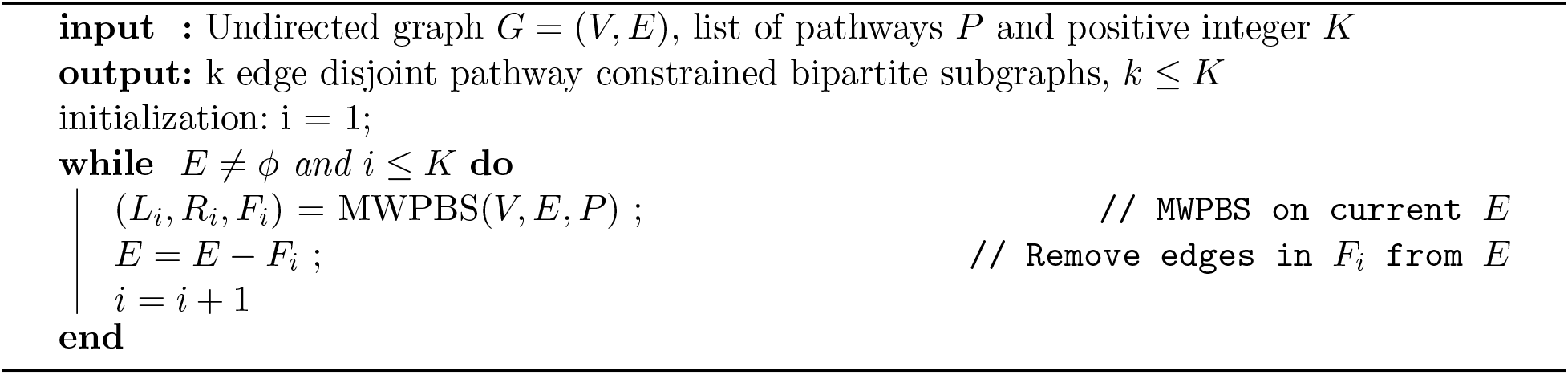
Greedy solution to the K-MWPBS problem

Note that the output bipartite sub-graphs of algorithm 1 can be concatenated to form a single subnetwork of the input *G*, as shown in figure 1.

Hypothetically, if the pathways considered are such that all the input genes are contained in every pathway, then the MWPBS problem is equivalent to solving the Maximum Weight Bipartite Subgraph Problem, which is NP-complete [21]. Therefore the MWPBS problem is NP-complete. Note that if each gene belongs to only one pathway, then the problem can be solved in polynomial time by ranking the weights of bipartite subgraphs formed by each pair of input pathways, and selecting the one with the highest weight.

In the following we formulate an Integer Linear Program (ILP) to solve the MWPBS problem. The advantage of an ILP is that it can leverage accurate and efficient heuristics, such as LP-based Branch and Bound and Cutting Plane methods, with readily available implementations in general-purpose solvers (e.g. Gurobi [22]) to find approximate solutions.

### 2.1 ILP Formulation for the MWPBS Problem

We assume all edge weights are positive and non-existent edges in the graph have weight 0 and *w*_*ij*_ = *w*_*ji*_ ∀*i, j*. Binary variables *e*_*ij*_ will be used to indicate the final selected edges in the bipartite subgraph. To obtain the maximum weighted subgraph we set the objective function to:

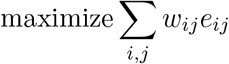

for every pair of nodes *i, j* such that *i* ≠ *j*. We now develop constraints that will ensure that the selected edges and their incident vertices form a bipartite subgraph. Define integer variable *x*_*i*_ for each vertex in the graph and integer variable *y*_*ij*_ for every pair of nodes *i, j* such that *i* ≠ *j*. We will restrict the ranges of these variables as follows:

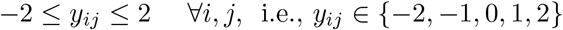

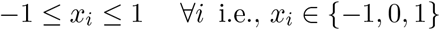

Variables *x*_*i*_ will be used to identify three subgroups of vertices of the input graph. Denote by *L* and *R* the left and right sets respectively of the selected bipartite graph, and by *O* all the remaining vertices in the input graph. Variables *x*_*i*_ = −1 will indicate vertices in *L*, *x*_*i*_ = 1 will indicate vertices in *R*, while *x*_*i*_ = 0 will indicate vertices in *O*.

For an edge in the required bipartite graph, i.e., between vertices *x*_*i*_ ∈ *L* and *x*_*j*_ ∈ *R*, we have *x*_*i*_ − *x*_*j*_ = −2 and for *x*_*i*_ ∈ *R* and *x*_*j*_ ∈ *L*, we have *x*_*i*_ − *x*_*j*_ = 2. For an edge between a vertex *x*_*i*_ ∈ *L* and a vertex *x*_*j*_ ∈ *O*, we have *x*_*i*_ − *x*_*j*_ = −1 and for *x*_*i*_ ∈ *O* and *x*_*j*_ ∈ *L*, we have *x*_*i*_ − *x*_*j*_ = 1. Similarly, for *x*_*i*_ ∈ *O* and *x*_*j*_ ∈ *R*, we have *x*_*i*_ − *x*_*j*_ = −1 and for *x*_*i*_ ∈ *R* and *x*_*j*_ ∈ *O*, we have *x*_*i*_ − *x*_*j*_ = 1. Finally, for *x*_*i*_, *x*_*j*_ in the same set (*O*, *L* or *R*), we have *x*_*i*_ − *x*_*j*_ = 0. So, we can identify the kind of edge using the following constraint:

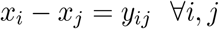

Note that for each pair of vertices, we shall obtain two values *y*_*ij*_ and *y*_*ji*_. Only one of them will affect the final objective function as described below.

Since we want to retain only the edges in the bipartite graph, i.e. where *y*_*ij*_ = 2, we will impose the constraints: if *e*_*ij*_ = 1 then *y*_*ij*_ ≥ 2 and if *e*_*ij*_ = 0 then *y*_*ij*_ ≤ 1. Since each edge is accounted for twice, once in each direction, excluding edges with *y*_*ij*_ = −2, shall not lead to missing any edges. These constraints can be imposed using an additional binary variable *z*_*ij*_ and a constant *M* ≥ 4 as follows:

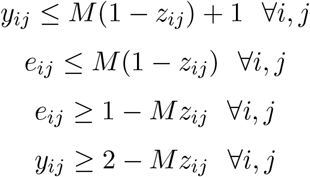

Note that when *z*_*ij*_ = 0 we have *e*_*ij*_ ≥ *1*, *y*_*ij*_ ≥ 2 (these edges are valid in the bipartite graph) and when *z*_*ij*_ = 1 we have *e*_*ij*_ ≤ 0, *y*_*ij*_ ≤ 1 (invalid edges for which *e*_*ij*_ are set to 0) as required.

#### Pathway Constraints

We now add constraints on the vertices to enforce each vertex set *L*, *R* in the bipartite subgraph to belong to a pathway. We first define binary variables *l*_*i*_ to indicate whether a node belongs to *L* or not: *l*_*i*_ = 1 iff *x*_*i*_ = −1. Similarly, we define binary variables to identify a vertex in *R*: *r*_*i*_ = 1 iff *x*_*i*_ = 1. These are set using the following constraints:

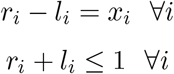

Let *N*_*p*_ be the number of pathways considered and *N*_*g*_ be the number of genes. Note that *N*_*g*_ = |*V*|. We assume a binary matrix *P* of dimension *N*_*g*_ × *N*_*p*_ indicating pathway membership where the *ij*^*th*^ cell is 1 if the *i*^*th*^ gene belongs to the *j*^*th*^ pathway, otherwise 0. Note that the summation ∑_*i*_ *P*_*it*_*l*_*i*_ for the *t*^*th*^ pathway (column in *P*) indicates the number of genes that are both in the *t*^*th*^ pathway and in the left set *L* of the bipartite graph. To ensure that every gene in the left set must belong to a pathway, we add the following constraints:

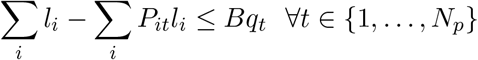

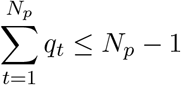

where *q*_*t*_ is a binary variable and *B* ≥ *N*_*g*_ is a large constant. At least one of the *q*_*t*_ must be set to 0 (otherwise, ∑_*t*_ *q*_*t*_ > *N*_*p*_ − 1) and this will make ∑_*i*_ *l*_*i*_ − ∑_*i*_*P*_*it*_*l*_*i*_ ≤ 0 ⇒ ∑_*i*_ *l*_*i*_ ≤ ∑_*i*_ *P*_*it*_*l*_*i*_. Since ∑_*i*_ *P*_*it*_*l*_*i*_ cannot be greater than ∑_*i*_ *l*_*i*_, this will set ∑_*i*_ *l*_*i*_ = ∑_*i*_*P*_*it*_*l*_*i*_ ensuring that all the genes in the left set are from the *t*^*th*^ pathway.

Similarly we ensure that the genes in the right set belong to a pathway:

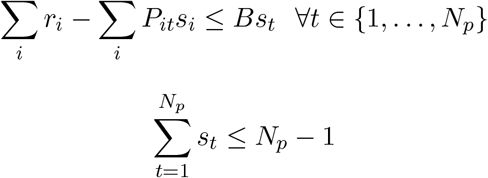

where *s*_*t*_ is a binary variable. Finally, every gene in *L* belongs to a pathway (say *P*_*l*_) and every gene in *R* belongs to a pathway (say *P*_*r*_), but we want the pathways *P*_*l*_, *P*_*r*_ to be different. This is done by the following constraint, which ensures that both *q*_*t*_, *s*_*t*_ are not set to 0 for the same *t*:

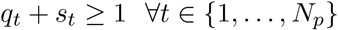

## 3 Experiments

### 3.1 Simulation Studies

We conduct simulation studies to evaluate the correctness of our algorithm. Since the MWPBS problem is NP-complete, optimal solutions cannot be found in polynomial time. We generate random networks under various settings and evaluate our ILP, both with and without pathway constraints, as well as our greedy algorithm for the K-MWPBS problem. Our results show that our ILP-based solutions can find optimal or near-optimal solutions as expected in the simulated conditions. Details are in Appendix A.

### 3.2 Real Data

#### Experiment Settings

We evaluate our algorithm on two graphs containing ME relations between genes in Breast cancer (BRCA) and Stomach cancer (STCA). ME relations were inferred using DISCOVER [8], separately for each cancer. DISCOVER infers ME using mutation data and accounts for heterogeneous alteration rates across tumor samples. All non-synonymous mutations (frameshift, missense, nonsense, nonstop and splice site) were used in our analysis. Mutation data for both the cancers were obtained from TCGA [50]. A total of 15,566 genes and 977 samples in BRCA and 16,920 genes and 393 tumor samples in STCA were used. 342 gene pairs in BRCA and 132 gene pairs in STCA were detected as ME (using p-value threshold < 0.05, with FDR-corrected p-value < 0.10 for STCA and 0.25 for BRCA respectively [9]), that were used to create the respective input graphs with negative logarithm of p-values as edge weights. We test our algorithm with signalling pathways from KEGG [29] and Hallmark [23]. A total of 169 KEGG pathways and 50 Hallmark pathways were used as input (Appendix B discusses how pathways were selected). We compute *between-pathway significance scores* by combining, using Fisher’s method [19], the p-values (obtained from DISCOVER) for all pairs of genes across the pathways. Table 4 in Appendix B shows the list of pathway name abbreviations used in our results.

### 3.3 Results

#### BRCA Dataset

Figure 2 shows the pathways selected by our algorithm and the ME relations between the pathways. There are 14 genes and 6 pathways in our solution among which seven are well-known cancer drivers: oncogenes (TP53, PIK3CA, MAP2K4, MAP3K1) and tumor-suppressor genes (TP53, PTEN, CDH1, NF1, MAP2K4, MAP3K1) [20]. Note that TP53 gene appears in both p53 and MAPK signaling pathways.

**Fig. 2.**
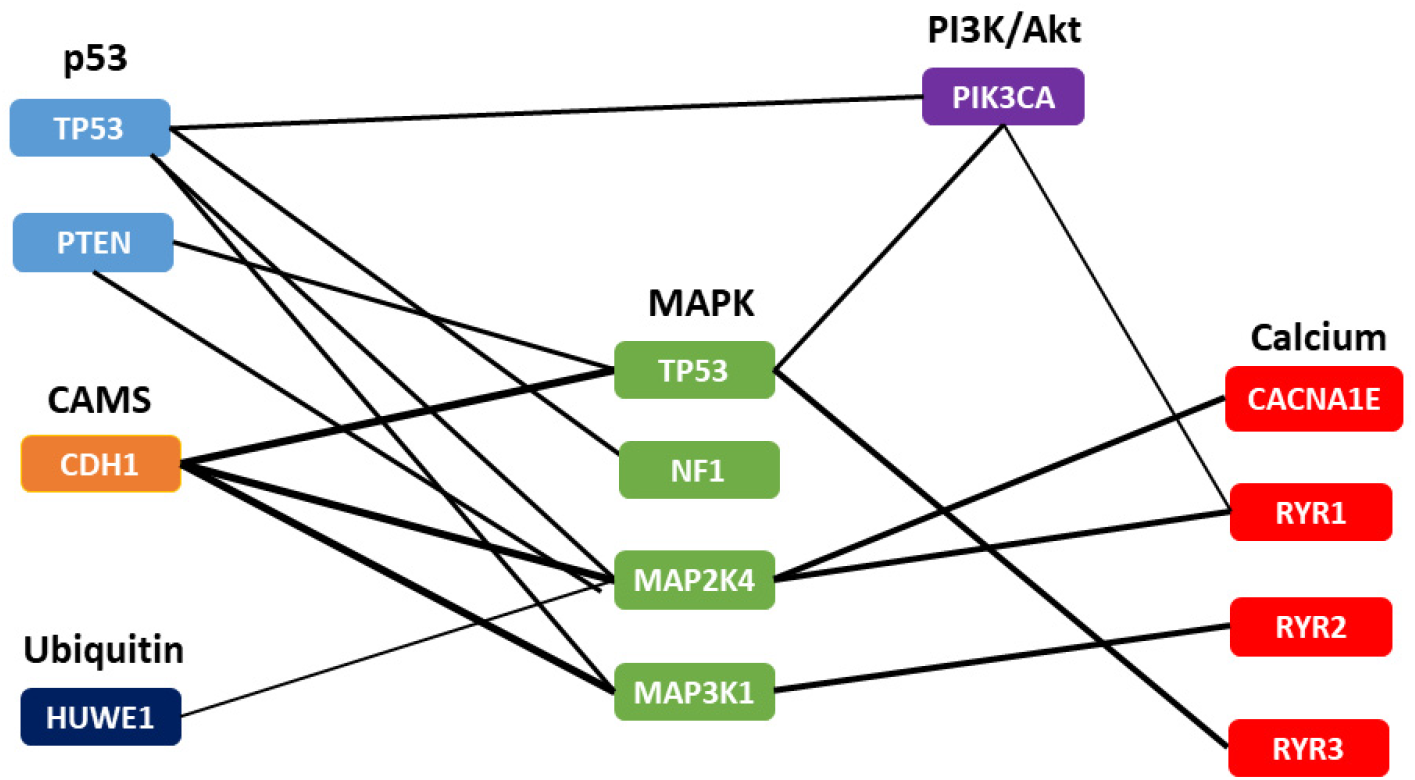
Pathways and genes selected by our method from the BRCA network. Nodes denote genes, edges indicate ME relations and each pathway is identified by a color. Thickness of edges is proportional to its significance.

The strongest ME relations between pathways was found between the CAMS and MAPK signaling pathways. It is known that CDH1 (or E-Cadherin) produces calcium-dependent cell-cell adhesion protein and is dependent on calcium ion (*Ca*^2+^) to function. Loss of function of this gene is thought to contribute to cancer progression by increasing proliferation, invasion and/or metastasis [52]. The second strongest ME relations among pathways was found between the MAPK and p53 signaling pathways. MAPK signaling pathway has been reported to regulate cell proliferation through a cross-talk of the MEK-ERK and JNK-JUN signaling pathways [60]. Cancers that have lost MAP3K1 or MAP2K4 fail to activate JNK-JUN signaling pathway [55], which in turn affects p53 activation. The activation of PI3K pathway via mutations in PIK3CA can activate p53 signaling in human cells, while activation of PIK3CA results in resistance to p53-related apoptosis in PTEN deficient cells [42]. Several studies elucidate the role of calcium and MAPK signaling in relation to cancer, particularly, breast cancer, e.g. [37]. Our algorithm mines the interactions among these pathways that are known to play important roles in breast cancer.

Cross-talks between the pathway pairs uncovered by our algorithm have been found useful in the context of breast cancer drug development. For instance, it has been reported that changes of intracellular *Ca*^2+^ activity could trigger numerous signaling pathways, in particular MAPK signaling pathway, that in turn perturb ER homeostasis resulting in ER stress-induced cell death, an association that has been used to design the cancer drug vemurafenib [16]. Calcium channel related genes (CACNA1E, RYR1, RYR2, RYR3) have featured in recent studies as potential drug targets [34, 2, 10, 38, 5]. In particular, calcium channel blockers such as Nifedipine and Verapamil, commonly used to treat hypertension, were found to be beneficial in some combinations of drugs [47]. Drug targeting the ubiquitin-proteasome pathway for cancer therapy has become an area of intense investigation [36]. Proteasome inhibitors effect apoptosis by decreased signaling through the MAPK pathway [36, 48]. The MAPK pathway itself is targeted by MAP2K4 (and MAP3K1) inhibiting drugs like Selumetinib and Trametinib [55]. The CAMS pathway has been found to be a useful target, e.g., through FDA-approved drugs like Docetaxel, in cases when cancer cells develop resistance to chemotherapeutic drugs [1].

An alteration map of 10 canonical signaling pathways from 33 cancer types in TCGA has been discussed in [39] (listed in Appendix B). We take the combination of the 6 pathways uncovered by our algorithm and these 10 therapeutically actionable canonical pathways selected in their study and compute their between-pathway significance scores, shown in figure 3. We note that the highest significance scores are for those pairs of pathways that our algorithm finds in figure 2, thereby providing additional empirical evidence of the correctness of our algorithm. We observe between-pathway ME interactions in breast cancer that were not reported in [39]. For instance, PI3K has low scores with the other canonical pathways (also seen in [39]) but has high scores with MAPK and Calcium pathways. Such cross-talks can be therapeutically exploited (as discussed in [39]) and dense pathway pairs with many ME interactions, uncovered by our algorithm, may be potential drug targets to be investigated further.

**Fig. 3.**
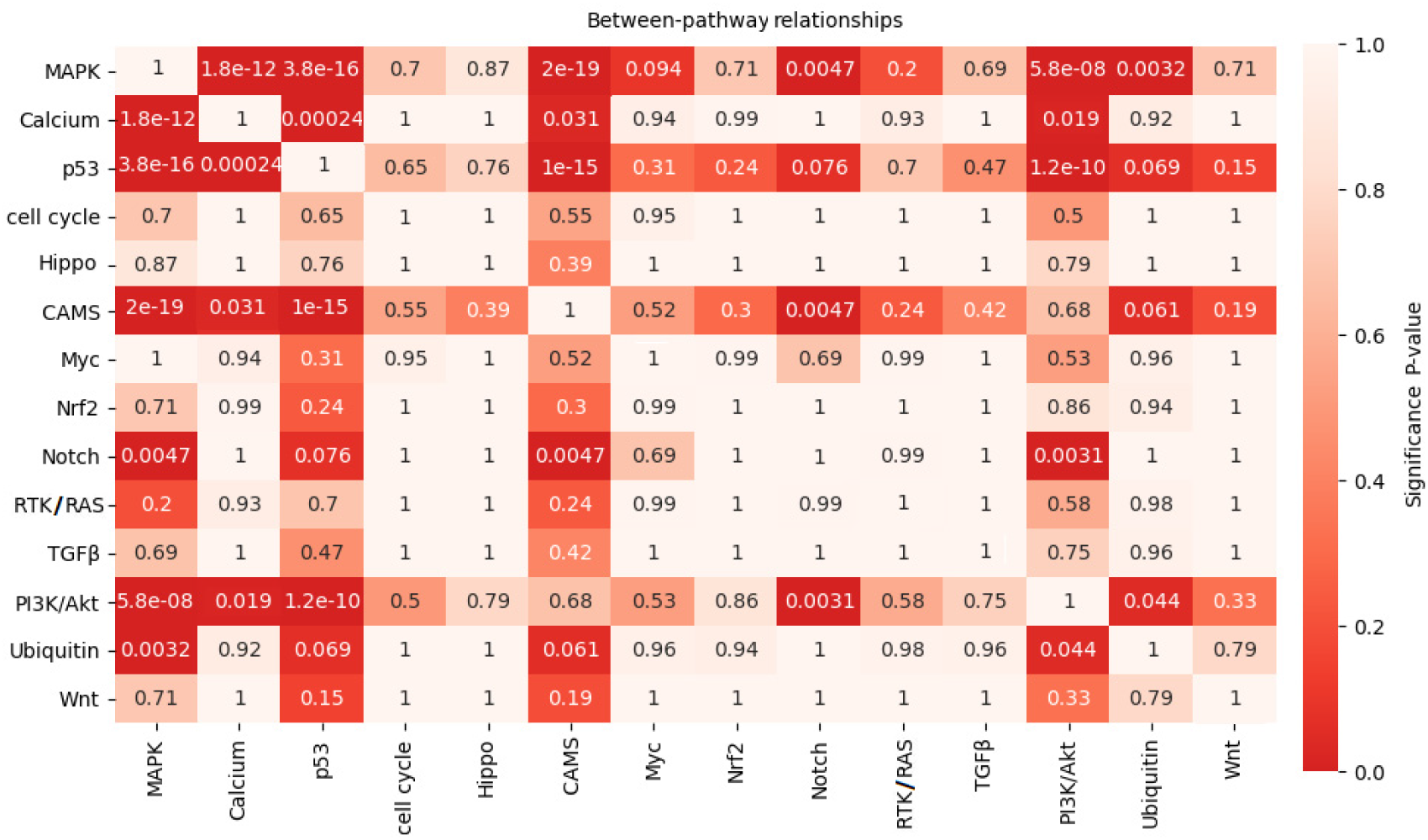
Between-pathway significance scores for pathways discovered by our method from the BRCA dataset along with 10 canonical signalling pathways in [39].

#### STCA Dataset

Figure 4 shows the pathways selected by our algorithm for the STCA dataset. There are 19 genes and 8 pathways in our solution. Seven of these are known driver genes: Oncogene (ERBB3, NOTCH1, TP53, PIK3CA) and tumor suppressor genes (MYH9, TP53, NOTCH1, CDH1, PTEN) [20]. Note that CDH1 gene appears in both CAMs and adherent junction pathways; PIK3CA gene appears in both apoptosis and focal adhesion pathways. Figure 5 shows the between-pathway significance scores for our pathways and the canonical signaling pathways [39]. We again observe that the highest significance scores are for pathway pairs obtained by our algorithm.

**Fig. 4.**
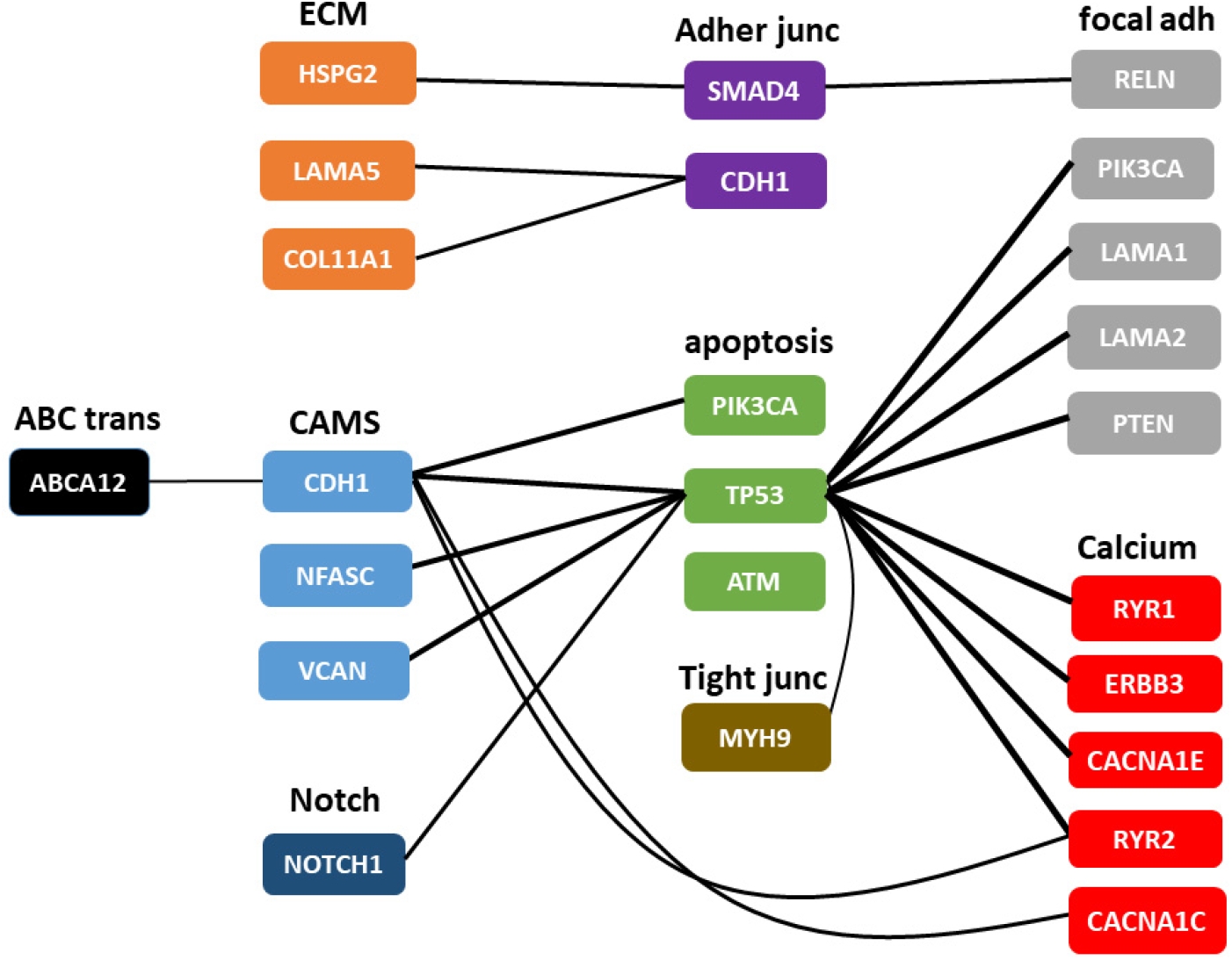
Pathways and genes selected by our method from the STCA network. Nodes denote genes, edges indicate ME relations and each pathway is identified by a color. Thickness of edges is proportional to its significance.

**Fig. 5.**
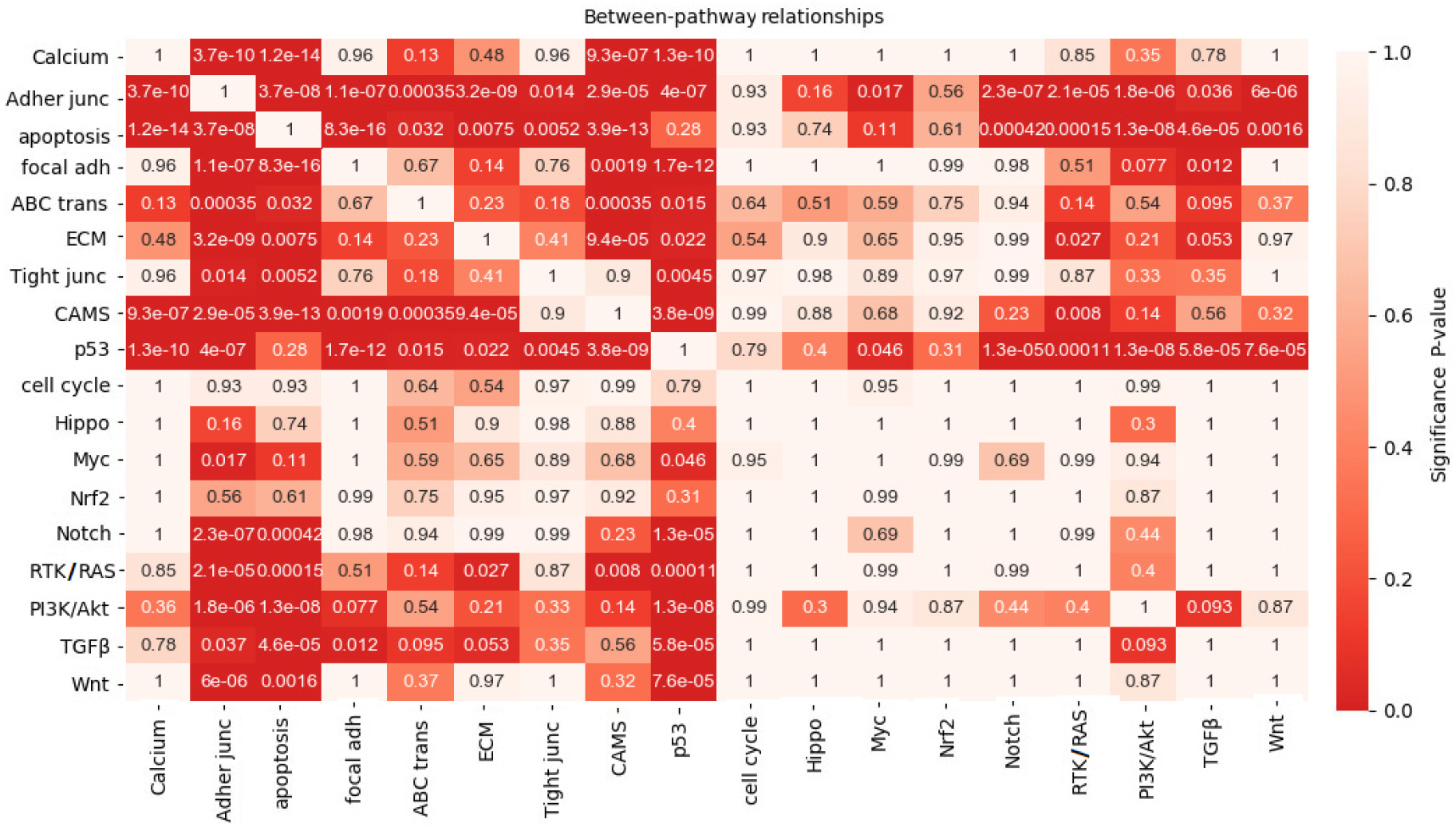
Between-pathway significance scores for pathways discovered by our method from the STCA dataset along with 10 canonical signalling pathways in [39].

Cross-talks between these pathways pairs have been reported in previous literature. The study in [44] discusses the links between CAMs/Tight junction(TJ)/Focal Adhesion (FA) proteins with apoptosis pathways, cell proliferation and inflammatory responses. NOTCH1 over-expression in the Notch signaling pathway is known to protect gastric carcinoma cells and reduce apoptosis [58]. The upregulation of IP3R3-mediated Ca2+ release through ER/mitochondrial cross-talk has been observed in gastric cancer cell lines; the depletion of ER Ca2+ stores itself is sufficient to cause ER stress and to induce apoptosis pathways [14]. FA provides mechanical linkage to ECM, and as a biochemical signaling hub to concentrate and direct numerous signaling proteins at sites of integrin binding and clustering [51]. Concurrently FA, ECM and adheren junctions actively contribute to the normal development, maintenance of the gastrointestinal tract and play important roles in stomach cancer [53] and all three pathways are mined by our algorithm.

These pathways, that are significantly involved in digestive carcinogenesis, have been used in targeted drug development. The drug Luteolin was found to suppress gastric cancer progression by suppression of the Notch signaling pathway [26]. The anti-inflammatory drug Ibuprofen is also useful as an adjuvant chemotherapy drug because of its role in downregulating Notch signaling [18]. The interaction between Notch and Apoptosis pathway is also targeted by the drug Penicilazaphilone C [49]. Several drugs, such as Rapamycin, Indomethacin and Cisplatin, target the apoptosis pathway [57, 11, 59]. Therapeutic targets for gastric cancer have also been found in Calcium and TJ pathways [41, 54, 61]. These studies show that pathway pairs highlighted by our algorithm have multiple targetable alterations that have been exploited in therapy development. The high-significance between-pathway interactions in figure 5 show multiple other potentially actionable pathways.

The pathway pairs given by our algorithm, and between-pathway significance scores for these pathways, when Hallmark pathways are used (instead of KEGG) are shown in Appendix C for both STCA and BRCA datasets.

## 4 Conclusion

We develop a new method to mine networks of mutual exclusivity interactions to find dense between-pathway associations. We formulate this as the problem of finding K edge-disjoint maximum weight pathway constrained bipartite subgraphs of the input network (the K-MWPBS problem). We develop an ILP-based formulation to solve the K-MWPBS problem. Unlike previous methods to find modules in networks our method allows input network nodes to be part of multiple subgraphs. This is an accurate model of the biological condition of genes participating in multiple pathways. Since these pathway pairs have multiple significant mutually exclusive interactions they are potential targets for novel drug development.

We empirically evaluate our method on two networks of mutual exclusivity interactions in breast and stomach cancer. We find that our method discovers pathways and their interactions that are known to play significant roles in cancer. These interactions have been independently discovered and studied earlier to gain deeper understanding of carcinogenesis. They have also found applications in targeted drug development. Our preliminary investigations thus illustrate the potential usefulness of our method in finding therapeutically actionable insights. The source code for our tool and experiments is available (at https://github.com/lianyh/K-MWPBS) for use by the community. Our method can be used to analyze networks of other interactions as well. We constrain our gene sets to belong to a pathway; however, other constraints can also be used (or added). Future studies can develop and analyze other such biologically relevant conditions.

## Appendix

## A Simulation Studies

We conduct simulations on synthetic networks to study the behavior of our ILP-based solution. The first simulation tests the ILP without pathway constraints. The second simulation tests the ILP with pathway constraints. Finally, the third simulation evaluates algorithm 1.

## Simulation 1

We generate random bipartite graphs with parameters *n, m* representing the number of nodes in the Left and Right sets respectively, and with the probability *p* of an edge across the sets. We then add *k* random edges to this graph, which may make it non-bipartite. Each edge had weight 1. We generate 4 sets of such networks with *p* = 0.5 and the parameter values given in table 1.

**Table 1.** Parameter values tested and results obtained in Simulation 1

| | $n$ | $m$ | $k$ | $\mu[e_S - e_G]$ | $\sigma[e_S - e_G]$ |
| --- | --- | --- | --- | --- | --- |
| I | 10 | 20 | 5 | 1.6 | 0.8 |
| II | 10 | 20 | 10 | 3.7 | 2.16 |
| III | 25 | 25 | 5 | 0 | 0 |
| IV | 25 | 25 | 10 | 0 | 0 |

For each set (I–IV) we generate 10 graphs. We then run the ILP without pathway constraints and evaluate the bipartite subgraph found. Let *H* be the input (simulated) graph to the ILP and let *S* be the solution returned by the ILP. Let *e*_*G*_ be the number of edges in simulated bipartite graph to which *k* random edges were added to form *H*. We know that *H* contains a bipartite subgraph with at least *e*_*G*_ edges. There may be more than *e*_*G*_ edges due to the added random edges. Let *e*_*S*_ be the number of edges in *S*.

We observe that the ILP correctly returns bipartite graphs in all the 40 solutions. Table 1 shows the mean (*μ*) and standard deviation (*σ*) of *e*_*S*_ − *e*_*G*_ over 10 runs for each set I–IV. In no simulation did *S* have lesser number of edges than *e*_*G*_. This suggests that the ILP found the maximum weighted bipartite subgraphs in each case for these relatively small networks.

## Simulation 2

We generate random bipartite graphs with parameters *n, m* representing the number of nodes in the Left and Right sets respectively, and with the probability *p* of an edge across the sets. Each of the *n* nodes in the Left set is assigned color red and each of the *m* nodes in the Right set is assigned color yellow, where a color denotes a distinct pathway. We add *j* random new nodes to the graph of colors red, yellow and black in equal proportions. We select *r* random nodes with replacement and randomly assign a new color (red or yellow or black) to the node, to have nodes with multiple colors. We then add *k* random edges to this graph, which may make it non-bipartite. Each edge had weight 1. We generate 4 sets of such networks with *p* = 0.5 and the parameter values given in table 2.

**Table 2.**
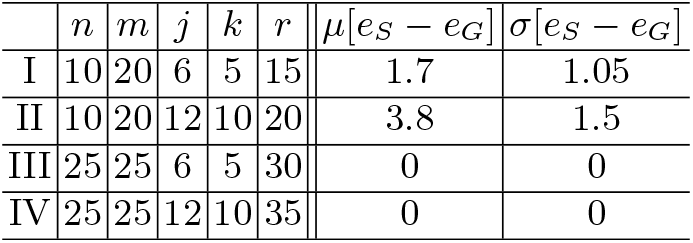
Parameter values tested and results obtained in Simulation 2

For each set (I–IV) we generate 10 graphs. We then run the ILP for the MWPBS problem and evaluate the bipartite subgraph found. The input list of pathways has 3 pathways, one for each color (red, yellow, black).

Let *H* be the input (simulated) graph to the ILP and let *S* be the solution returned by the ILP. Let *e*_*G*_ be the number of edges in simulated bipartite graph to which *k* random edges were added to form *H*. We know that *H* contains a bipartite subgraph with at least *e*_*G*_ edges. There may be more than *e*_*G*_ edges due to the added random edges. Let *e*_*S*_ be the number of edges in *S*.

We observe that the ILP correctly returns bipartite graphs in all the 40 solutions over simulation sets I–IV, and all the nodes in the left/right node sets are of one color, with the two colors being different. Table 1 shows the mean (*μ*) and standard deviation (*σ*) of *e*_*S*_ − *e*_*G*_ over 10 runs for each set I–IV. In no simulation did *S* have lesser number of edges than *e*_*G*_. This suggests that the ILP found the maximum weighted bipartite subgraphs in each case for these relatively small networks.

## Simulation 3

We generate 6 sets of nodes where each set contains 10 nodes and has a distinct color. We generate 6 bipartite graphs with number of edges 100, 50, 25, 15, 10, 5 by randomly selecting pairs of node sets. The same set may be selected more than once but the same pair of node sets is not selected more than once. These graphs are merged into one graph. We add *j* random new nodes to the graph where one of the six colors is randomly assigned to each added node. We select *r* = 30 random nodes with replacement and randomly assign a new color (from the six colors) to the node, to have nodes with multiple colors. We then add *k* random edges to this graph. Each edge has weight 1. We generate 4 sets of such networks with the parameter values given in table 3. For each set (I–IV) we generate 10 graphs. We then run our algorithm 1 for the K-MWPBS problem, with *K* = 6 and evaluate the bipartite subgraphs found. The input list of pathways has 6 pathways, one for each color used.

**Table 3.** Parameter values tested and results obtained in Simulation 3

| | $j$ | $k$ | $\mu[E_S - E_G]$ | $\sigma[E_S - E_G]$ |
| --- | --- | --- | --- | --- |
| I | 20 | 10 | 20 | 0 |
| II | 40 | 20 | 39.5 | 0.7 |
| III | 30 | 10 | 30 | 0 |
| IV | 60 | 20 | 19.4 | 4.25 |

Let *H* be the input graph to the algorithm and let *S*_1_ − −*S*_6_ be the 6 bipartite graphs returned by the ILP. Let *E*_*G*_ be the total number of edges in all the bipartite graphs merged to which *k* random edges were added to form *H*. We know that *H* contains 6 bipartite subgraphs with at least *E*_*G*_ edges. There may be more than *E*_*G*_ edges due to the added random edges. Let *E*_*S*_ be the total number of edges in *S*_1_ − −*S*_6_.

We observe that our algorithm correctly returns bipartite subgraphs in all the 40 solutions over simulation sets I–IV, and all the nodes in the left/right node sets, in each subgraph, are of one color, with the two colors being different. The number of bipartite subgraphs may be less than six as larger bipartite subgraphs may be formed due to random addition of colors and edges in the simulation. Table 3 shows the mean (*μ*) and standard deviation (*σ*) of *E*_*S*_ − *E*_*G*_ over 10 runs for each set I–IV. In no simulation was *E*_*S*_ less than *E*_*G*_. This suggests that our algorithm found six maximum weighted bipartite subgraphs in each case for these relatively small networks.

## B Experiment Settings

To focus on “signaling pathways”, we excluded the following 17 pathways in KEGG: huntingtons disease, pathways in cancer, colorectal cancer, pancreatic cancer, endometrial cancer, prostrate cancer, thyroid cancer, melanoma, bladder cancer, chronic myeloid leukemia, acute myloid leukemia, small cell lung cancer, non small cell lung cancer, hypertrophic cardiomyopathy HCM, arrhythmogenic right ventricular cardiomyopathy arvc, dilated cardiomyopathy, viral myocarditis. From Hallmark we do not exclude any pathways but utilize all existing 50 pathways.

Table 4 shows the list of pathway name abbreviations used in our figures.

**Table 4.** Pathway name abbreviations used in figures.

| Pathway abbreviations | Full Pathway names |
| --- | --- |
| MAPK | MAPK Signaling Pathway |
| Calcium | Calcium Signaling Pathway |
| p53 | p53 Signaling Pathway |
| cell cycle | Cell Cycle Pathway |
| Hippo | Hippo Signaling Pathway |
| CAMS | Cell adhesion molecules Pathway |
| Myc | Myc Pathways |
| Nrf2 | NRF2 Pathway |
| RTK/RAS | RTK-RAS Signaling Pathway |
| TGFβ | TGF Beta Signaling Pathway |
| PI3K/Akt | PI3K-Akt Pathway |
| Ubiquitin | Ubiquitin Mediated Proteolysis Pathway |
| Wnt | Wnt Signaling Pathway |
| Adher junc | Adherens Junction Pathway |
| apoptosis | Apoptosis Pathway |
| focal adh | Focal Adhesion Pathway |
| ECM | ECM-receptor interaction Pathway |
| ABC trans | ABC Transporter Pathway |
| Tight junc | Tight Junction Pathway |
| Notch | Notch Signaling Pathway |
| H p53 | Hallmark p53 Pathway |
| H Apical junc | Hallmark Apical Junction Pathway |
| H Compl | Hallmark Complement Pathway |
| H EMT | Hallmark Epithelial-mesenchymal Transition Pathway |
| H IL2 STAT5 | Hallmark IL2 STAT5 Signaling Pathway |
| H Myogenesis | Hallmark Myogenesis Pathway |
| H KRAS s UP | Hallmark KRAS Signaling UP Pathway |
| H KRAS s DN | Hallmark KRAS Signaling DN Pathway |
| H UV R DN | Hallmark UV Response DN Pathway |
| H UV R UP | Hallmark UV Response UP Pathway |

The 10 canonical signaling pathways discussed in [39] are:

1. Cell Cycle Pathway
2. Hippo Signaling Pathway
3. Myc Pathway
4. NRF2 Pathway
5. RTK-RAS Signaling Pathway
6. TGF Beta Signaling Pathway
7. PI3K-Akt Pathway
8. Notch Signaling Pathway
9. p53 Signaling Pathway
10. Wnt Signaling Pathway

## C Analysis with Hallmark pathways

The pathway pairs given by our algorithm when Hallmark pathways are used are shown in figure 6 for BRCA and in figure 8 for STCA. Between-pathway significance scores for these pathways and the canonical signaling pathways are shown in figure 7 for BRCA and figure 9 for STCA.

**Fig. 6.**
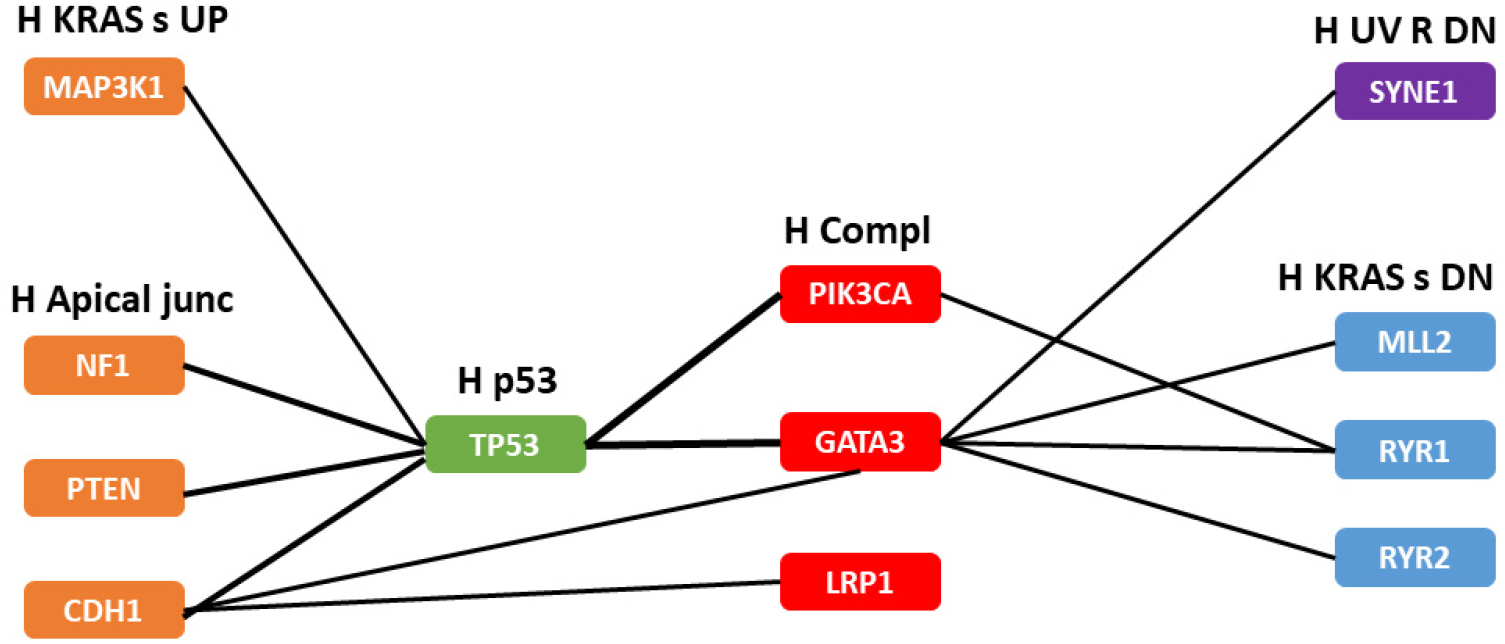
Pathways and genes selected by our method from the BRCA network. Nodes denote genes, edges indicate ME relations and each pathway is identified by a color. Thickness of edges is proportional to its significance.

**Fig. 7.**
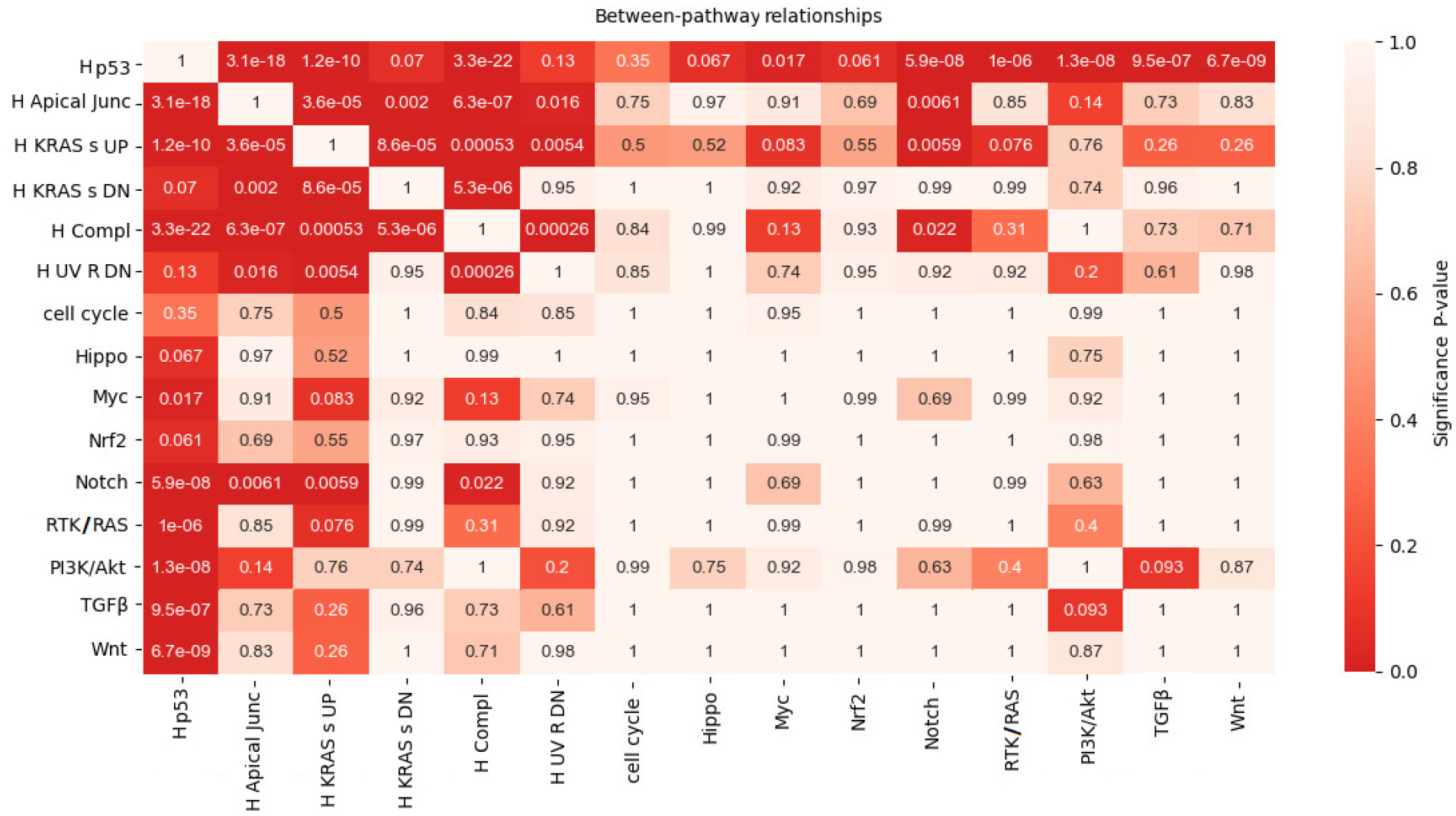
Between-pathway significance scores for pathways discovered by our method from the BRCA dataset along with 10 canonical signalling pathways in [39].

**Fig. 8.**
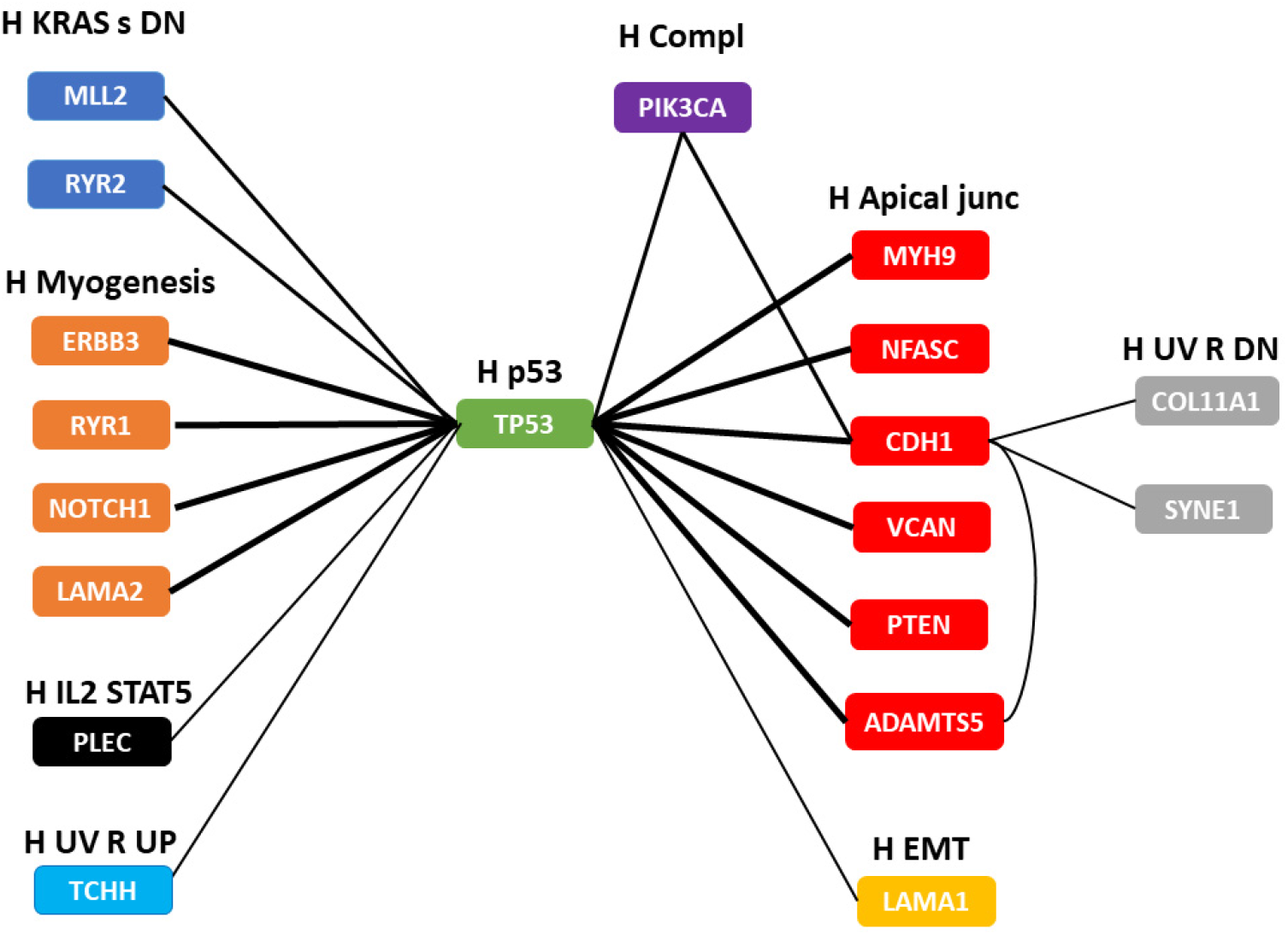
Pathways and genes selected by our method from the STCA network. Nodes denote genes, edges indicate ME relations and each pathway is identified by a color. Thickness of edges is proportional to its significance.

**Fig. 9.**
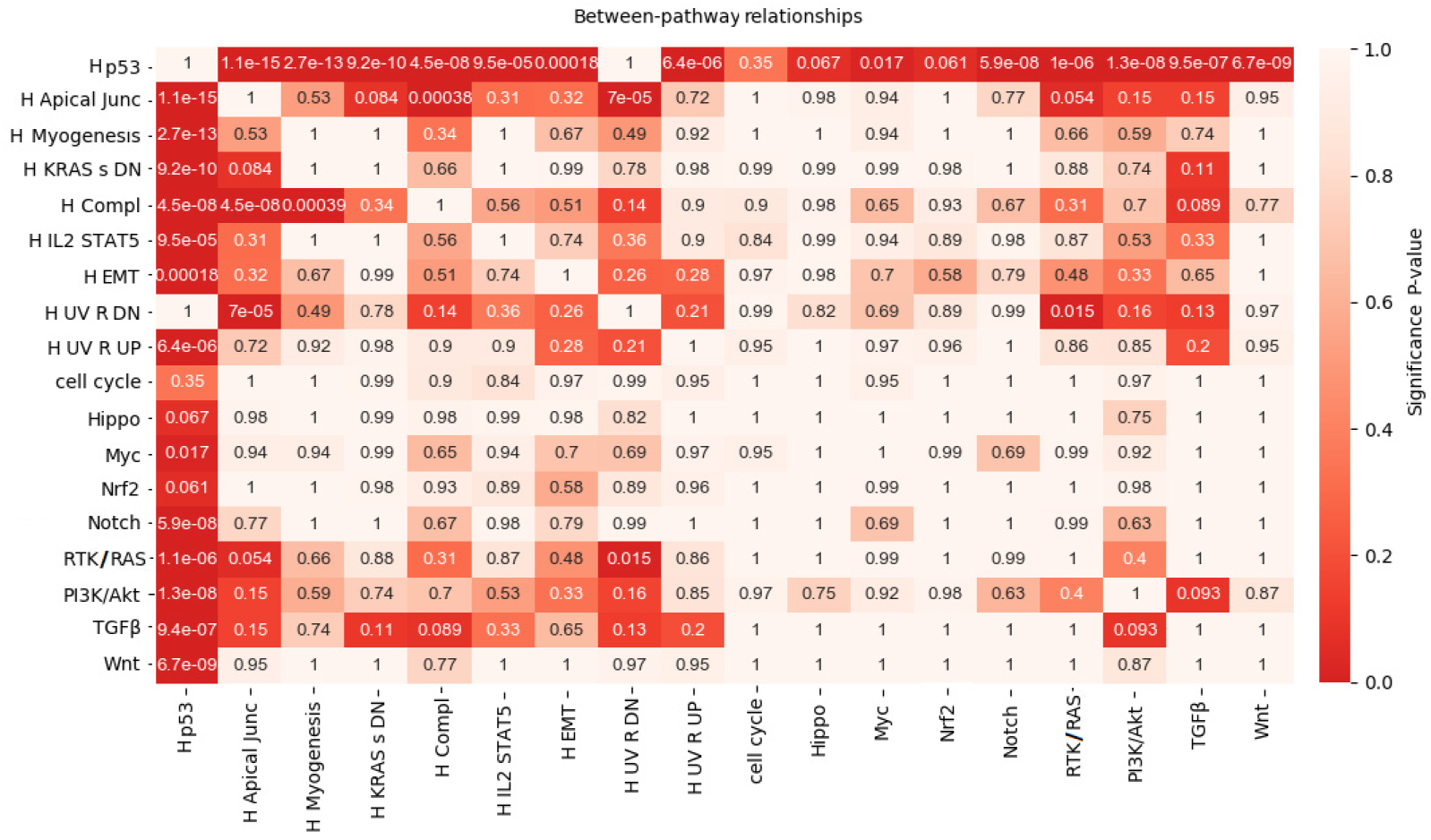
Between-pathway significance scores for pathways discovered by our method from the STCA dataset along with 10 canonical signalling pathways in [39].

The common genes identified in BRCA data by our method in both KEGG and Hallmark pathways are TP53, PIK3CA, MAP3K1, PTEN, CDH1, NF1, RYR1, and RYR2. Some genes such as GATA3, SYNE1 only exist in Hallmark whereas CACNA1E, MAP2K4 genes only exist in KEGG.

In the STCA dataset, common genes identified by our method in both KEGG and Hallmark are TP53, PIK3CA, CDH1, NOTCH1, PTEN, MYH9, ERBB3, NFASC, COL11A1, LAMA2, and VCAN. Some other identified genes,SYNE1, ADAMTS5, MLL2, PLEC only exist in Hallmark and CACNA1E, CACNA1C, ABCA12, SMAD4 genes only exist in KEGG.

